# Rapid Isolation and Genomic Characterization of Sewage-Derived Bacteriophages Targeting Multidrug-Resistant Avian Pathogenic Escherichia coli in Poultry

**DOI:** 10.64898/2026.02.13.705853

**Authors:** Rajindra Napit, Anupama Gurung, Ashok Chaudhary, Machchendra Thapa, Roji Raut, Ajit Poudel, Sajisha Prajapati, Dhiraj Puri, Dibesh Karmacharya

**Affiliations:** Center for Molecular Dynamics Nepal, Thapathali, Nepal; Central Department of Biotechnology, Tribhuvan University, Kirtipur, Nepal; University of Queensland, Queensland, Australia; BIOVAC Nepal Pvt. Ltd., Kathmandu, Nepal

**Keywords:** Bacteriophage therapy, antimicrobial resistance, avian pathogenic *E. coli*, phage genomics, sewage-derived phages

## Abstract

Antimicrobial resistance (AMR) in *Escherichia coli* is a major threat to global poultry production, particularly for controlling colibacillosis caused by avian pathogenic *E. coli* (APEC). Intensive antibiotic use has accelerated the emergence of multidrug-resistant (MDR) strains, undermining treatment efficacy and facilitating zoonotic transmission of resistance genes. This study evaluated bacteriophage therapy as a targeted, antibiotic-independent strategy to control MDR *E. coli* in Nepalese poultry systems.

Seventeen *E. coli* isolates were obtained from commercial broiler and local-breed chicken farms. Using the double-layer agar method, 18 lytic phages were isolated from three urban sewage samples, infecting eight isolates. Real Time PCR screening revealed extensive AMR profiles in six isolates; one broiler-derived strain (EcI8) harbored 30 resistance genes, including carbapenemases, ESBLs, aminoglycoside, quinolone, and last-resort antibiotic determinants associated with class 1 and 3 integrons.

Purified phages were whole-genome sequenced and all of them belonged to the genera *Tequatrovirus*, *Phapecoctavirus*, *Vequintavirus*, and *Gamaleyavirus*—lineages previously used in veterinary and human phage therapy. Several exhibited broad host ranges (up to seven isolates) and lacked detectable virulence or resistance genes; they also encoded potent lytic enzymes, including endolysins, holins, spanins, and transglycosylases. Notably, one *Phapecoctavirus* (PG7) showed close similarity to established APEC-infecting phages.

These findings demonstrate that sewage can be a readily accessible source of safe, therapeutically relevant phages targeting highly resistant poultry *E. coli*. Phage cocktails and derived enzybiotics delivered through drinking water or feed offer a scalable alternative for preventing and treating colibacillosis, reducing antibiotic dependence, and mitigating zoonotic AMR risks. This study establishes a rapid, reproducible pipeline for local phage isolation and genomic validation, supporting field deployment of tailored phage-based interventions in resource-limited poultry systems.

## Introduction

Antimicrobial resistance (AMR) is a growing concern in the poultry industry, and *Escherichia coli* (*E .coli*) is one of the most prevalent bacterial species showing high levels of resistance to several antibiotics, including tetracycline, ampicillin, and sulphonamides [1]. *E. coli,* a bacterium that is a common inhabitant of the gastrointestinal tract of vertebrates and warm-blooded animals, is the most prevalent commensal bacteria which can cause a variety of infections [2]. Among them, Avian pathogenic *E.coli* (APEC), is a major causative agent of respiratory, systemic, and reproductive disease in poultry worldwide [3–5]. Antimicrobials are being used in the treatment of colibacillosis in poultry which has been an important tool in increasing productivity and reducing mortality and the economic burdens in poultry [6]; however, the emergence of antibiotic-resistant strains of *E .coli* poses a significant threat to the efficacy of antibiotic treatment [7].

Bacteriophages are viruses that specifically infect bacteria, attaching to their hosts and injecting their genetic material to hijack the bacterial machinery for replication [8]. Phages propagate via two main life cycles: the lytic cycle, in which the phage replicates and ultimately lyses (kills) the host cell, and the lysogenic cycle, where the phage genome integrates into the host chromosome and is replicated passively as a prophage over many generations [9]. Under certain stress conditions, the prophage can be induced to enter the lytic cycle, resulting in bacterial cell lysis [10]. Conversion of lytic phage into lysogenic phage is also guided by molecular phage quorum sensing based on the concentration of arbitrium molecules as an increase in phage number leads to an increase in arbitrium molecules concentration which switches lytic phage to lysogenic phage [11].

Since Bacteriophages discovery by William Twort in 1915 and their potential to kill bacteria by Felix d’ Herelle in 1917 [12], bacteriophage research as a therapeutic agent has been continuing [13]. But its significance was minimized after antibiotics were discovered in 1945 and subsequently used in treating bacterial infections [14].

According to the GLASS Report 2022, antimicrobial consumption was highest in the African, South-East Asian, and Western Pacific regions, with a median use of 16.6 defined daily doses (DDD) per 1,000 inhabitants per day. The median proportional consumption comprised 67% Access group antibiotics and 31% Watch group antibiotics [15]. In 2018, Nepal consumed 357.8 tons of antibacterial, with 74% belonging to the Watch group and only 26% to the Access group, a pattern observed in both clinical and poultry settings that substantially elevates the risk of antimicrobial resistance in the country [16].

Bacteriophage (also referred to as Phage) therapy has emerged as a promising alternative to antibiotics in combating bacterial infections, particularly in the context of rising antimicrobial resistance (AMR) [17]. However, several critical drawbacks limit its widespread clinical application and necessitate innovative approaches for deployment. Phages may exhibit lysogenic behavior, integrating into the host genome and potentially transferring virulence or resistance genes [18,19]. Host immune responses can neutralize phage particles, reducing therapeutic efficacy and complicating systemic administration [19–21]. Inherently narrow host ranges require extensive phage collections or cocktails to ensure coverage [18,22]. Most significantly, bacterial populations can rapidly evolve resistance to therapeutic phages—often within days of treatment initiation—rendering initially effective phages obsolete and potentially undermining preapproved or static phage libraries [18,23,24]. This rapid resistance evolution makes the classical approach of extensive *ex ante* phage isolation, comprehensive characterization, regulatory approval, banking, and subsequent deployment impractical for dynamic clinical scenarios where pathogen genotypes are continuously evolving.

This study presents a rapid-response strategy to mitigate colibacillosis and antimicrobial resistance (AMR) in avian pathogenic *Escherichia coli* (APEC). Recent surveillance indicates alarmingly high resistance rates among APEC isolates, including >98% resistance to ampicillin, multidrug resistance in over 90% of strains, and emerging colistin resistance (*mcr-1*) in more than half of tested isolates from poultry samples (23,24). These infections impose substantial economic burdens on the poultry industry, resulting in multi-million-dollar annual losses due to mortality, reduced productivity, and treatment costs [8,10]. Although bacteriophages are promising alternatives to antibiotics in veterinary medicine, their deployment has been constrained by concerns related to lysogenicity, host specificity, immunological responses, and time-intensive characterization workflows (13,14,17,18).

To address these limitations, we implemented a rapid-response framework that combines accelerated phage isolation using the double-layer agar method with streamlined genomic characterization via Nextera XT library preparation and Illumina sequencing. Designed as an outbreak management tool, this approach enables a 4–7-day turnaround from pathogen isolation to strain-specific, genomically validated phage deployment at the flock level. By rapidly screening phages for lysogeny markers, virulence factors, and antimicrobial resistance genes, this strategy overcomes the inherent lag of static phage libraries in the face of rapidly evolving bacterial populations. By integrating classical phage isolation with contemporary genomic validation, our pipeline provides an adaptive and scalable approach for generating therapeutically relevant, strain-specific phages, enhancing the practical feasibility of phage therapy for controlling emerging AMR in poultry production systems.

## Material and methods

This study used an integrated microbiological and genomic approach to isolate and characterize *Escherichia coli* and associated bacteriophages from poultry farms in Nepal. Biological and environmental samples were collected from nine farms across five districts, along with urban sewage samples for phage isolation. *E. coli* isolates were identified using selective culture, biochemical testing, and 16S rRNA gene sequencing. Host strains were used for phage enrichment, plaque isolation, purification, and cross-infectivity testing to assess lytic activity and host range. Phage-susceptible *E. coli* isolates were screened for antimicrobial resistance genes using high-throughput PCR. Purified phages were concentrated by polyethylene glycol precipitation, followed by whole-genome sequencing on the Illumina MiSeq platform. Bioinformatic analyses included quality control, de novo assembly, viral genome validation, taxonomic classification, and screening for antimicrobial resistance and virulence genes (Figure 1).

**Figure 1:**
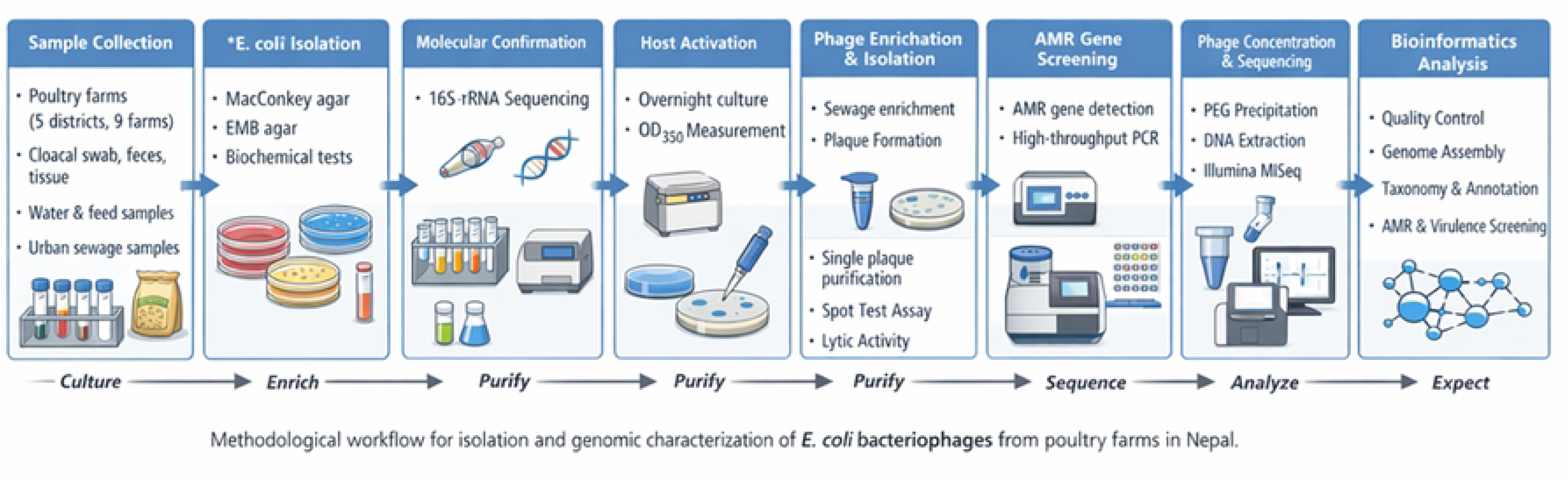
Overview of the methodological workflow used for isolation, purification, genomic sequencing, and bioinformatic analysis of Escherichia coli bacteriophages recovered from poultry farms and urban sewage samples in Nepal.

### Sample Collection

Biological samples—including cloacal swabs, feces, oral swabs, tissues, environmental water, and feed—were collected from poultry farms across Nepal (Kavre, Chitwan, Tanahun, Dhading, and Nuwakot) for the isolation of *Escherichia coli* from March to June, 2022. Samples were obtained from nine farms across five districts, with two independent sampling sets collected per farm (Figure 2). The sampled populations represented four poultry types: local breeds (endemic), commercial layers, commercial broilers, and mixed breeds. All samples were collected in sterile 2 ml cryovials containing 500 µl of 50% glycerol, maintained under cold-chain conditions, and transported to the laboratory for further processing. For bacteriophage isolation and recovery, 50 ml sewage grab samples were collected in sterile 50 ml Falcon tubes from three locations in the Kathmandu Valley (Balaju, Dallu and Thapathali).

**Figure 2:**
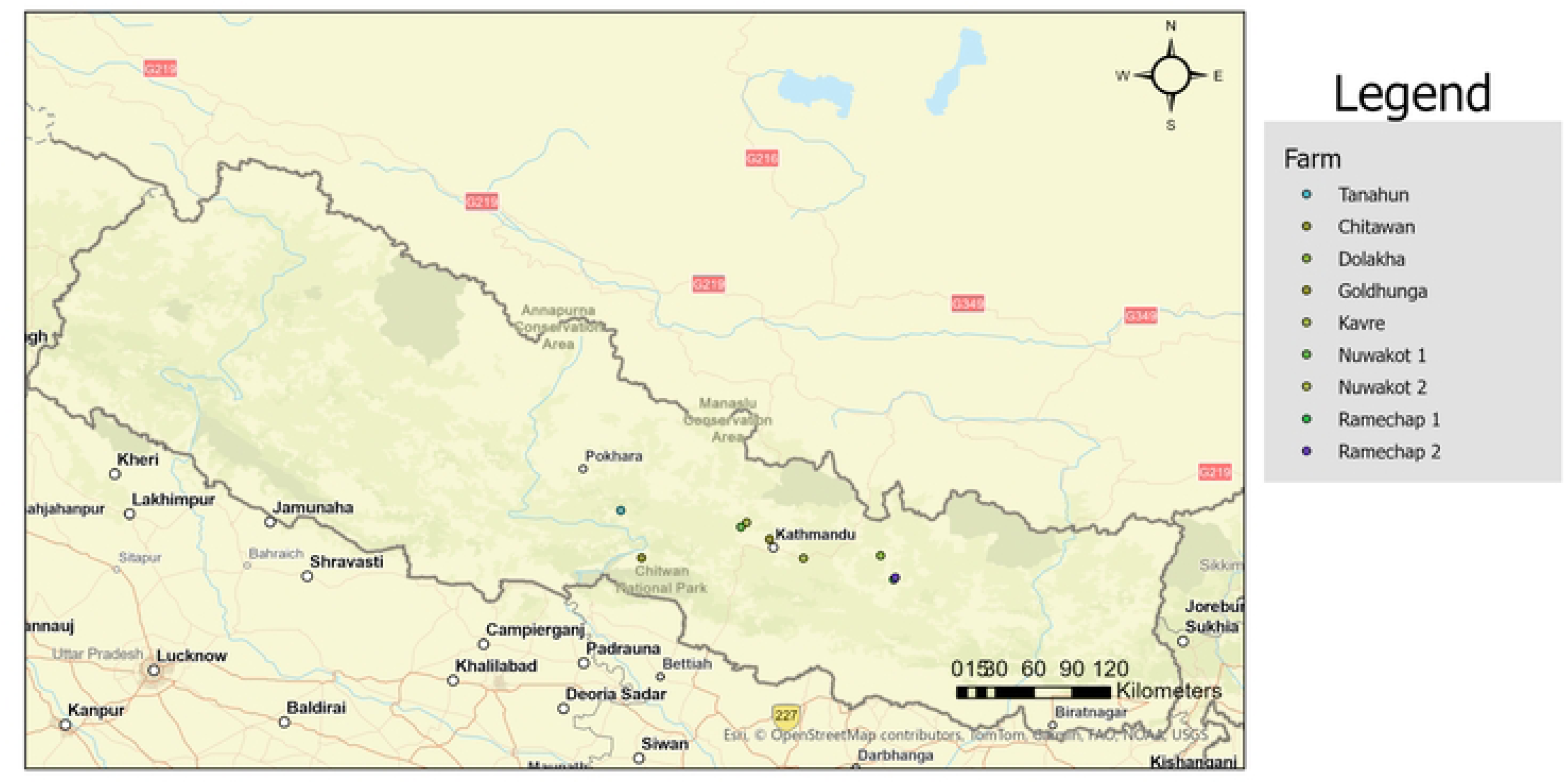
Geographic distribution of poultry farms sampled across Nepal. Sampling sites span hill and lowland regions, including Ramechhap, Dolakha, Kavre, Nuwakot, Tanahun, Chitwan, and Goldhunga (Kathmandu Valley). At all sites, host-associated (cloacal swab, oral swab, faeces) and environmental (farm water, farm feed) samples were collected, while gizzard and proventriculus tissues were obtained only at the Goldhunga site. The map illustrates the spatial coverage of poultry health and environmental surveillance.

### Screening for Escherichia coli (E. coli)

All the samples from various poultry farms were streaked on a bacterial culture plate with MacConkey Agar (Hi-Media, M082-500G) and incubated at 37°C for 24 h. The lactose fermenting colonies were selected and enriched in nutrient broth and sub-cultured with Eosin Methylene Blue Agar (Hi-Media, M317-500G). Sub-cultured plates were incubated at 37 °C for 24 h. After incubation, colonies exhibiting a characteristic metallic green sheen were presumptively identified as *Escherichia coli*. These isolates were further subjected to biochemical characterization, including catalase, oxidase, triple sugar iron agar (TSIA), sulfide–indole–motility (SIM), methyl red–Voges–Proskauer (MR–VP), citrate utilization, and urease tests (HiMedia, India), following standard protocols [25]. Molecular characterization was subsequently performed to confirm the identity of *E. coli* isolates.

### 16S rRNA Sequencing for *E. coli*

The isolated putative colonies of *E. coli* were further confirmed with 16S amplicon DNA sequence validation. The identified *E. coli* colonies were then activated in nutrient broth overnight and concentrated to obtain the bacterial pellet. The bacterial pellet was used for genomic DNA extraction using the QIAamp Fast DNA Stool Mini Kit (Qiagen; Cat. No. 51604), following the manufacturer’s instructions. Extracted DNA was subjected to amplification of the 16S rRNA gene using the universal bacterial primers 16sBakt341F and 16sBakt805R. PCR amplification was carried out in a 25 µL reaction volume containing 5 µL each of forward and reverse primers (1 pmol/µL), 12.5 µL of 2× KAPA HiFi HotStart ReadyMix (Roche, Switzerland), and 2.5 µL of template DNA. Thermocycling conditions consisted of an initial denaturation at 95 °C for 3 min, followed by 35 cycles of denaturation at 98 °C for 30 s, annealing at 65 °C for 25 s, and extension at 72 °C for 20 s, with a final extension at 72 °C for 5 min. PCR amplicons were resolved on a 1.5% agarose gel stained with GelRed and visualized after electrophoresis at 90 V for 90 min. The expected amplicon size (∼600 bp) was confirmed using a 100 bp DNA ladder (Solis BioDyne). The amplified product was purified with 1X AMPure XP beads (Beckman Coulter), and library preparation, including tagmentation and indexing, was carried out using the Nextera XT kit. The prepared library was then sequenced on the Illumina MiSeq platform.

### Activation of *E. coli*

*E. coli* activation was performed by inoculating 2 mL of nutrient broth (HiMedia; M002) and incubating the culture overnight at 37 °C with shaking at 100 rpm. Bacterial growth was assessed the following day by measuring optical density at 630 nm using a BioTek microplate reader to estimate growth potential and host cell density.

### Isolation of Phages

Phage enrichment was performed prior to plaque isolation using differential centrifugation of sewage samples. Briefly, sewage samples were centrifuged at 1,000 rpm for 1 min to remove coarse debris. The supernatant was then transferred to a sterile 50 mL Falcon tube and centrifuged at 8,000 rpm for 5 min to remove bacterial cells. The clarified supernatant was enriched by mixing with an equal volume of tryptone soya broth (TSB; HiMedia, M011) supplemented with 10 mM MgCl_₂_, as previously described [26]. Overnight-activated *E. coli* cultures were adjusted to an optical density (OD) of 0.1 and added to the enrichment mixture. The enrichment culture was incubated at 37 °C for 5 h with shaking to promote phage amplification.

The enriched broth was centrifuged at 2000rpm for 2min, and the supernatant was transferred to a fresh tube and centrifuged at 8000rpm for 5 minutes. Further serial dilution of the enriched broth was performed from 10^-1^ to 10^-6^. Soft agar overlay method (0.7% agar-agar) was used for the plaque assay of phages [27,28] using 10^-4^ and 10^-6^ dilution against 1% host. Plates were incubated at 37°C for 24 h and development of plaques were observed.

### AMR genes detection in *E. coli*

*E. coli* isolates that were susceptible to phage infection were subsequently screened for antimicrobial resistance (AMR) genes in their genomes. AMR genes were detected in the isolates using qualitative PCR (Resistomap, Finland) performed in triplicate. Each isolate was tested for over 100 different AMR genes.

### Purification of Phages

Phage purification was carried out by selecting individual clear plaques based on size and morphological uniformity. Single plaques were aseptically excised from the agar surface using sterile filter tips and enriched in tryptic soy broth (TSB; HiMedia) supplemented with 10 mM MgCl_₂_, followed by incubation at 37 °C for 5 h. Serial dilutions of the enriched phage suspension were subjected to spot tests on host bacterial lawns. A pour plate assay using an appropriate dilution was subsequently performed to assess plaque uniformity. Phage purity was confirmed through multiple successive passages of single plaques exhibiting consistent morphology [27,28].

### Cross infectivity of phage isolates

Spot test technique was performed for cross infectivity of phage isolates to determine the lytic activity of each phage isolates. All the isolated phages were enriched with different host in TSB media incubating for 5 h at 37°C. Top agar overlay of each host was prepared on tryptic soy agar (TSA) and left to solidify. Ten microliters of each enriched phages were spotted on the solidified soft agar and allowed to air dry on laminar air flow [29]. The plates were incubated at 37°C for 24 h. The plates were observed for presence of plaques after incubation. Phage cross-infectivity was assessed based on three lysis phenotypes: complete lysis, partial lysis, and turbid lysis, as previously described.

### Phage Precipitation by Polyethylene Glycol (PEG 6000)

Twenty milliliters of propagated, purified phage lysate were mixed with an equal volume of 30% polyethylene glycol (PEG) containing 2% (w/v) NaCl [30]. The mixture was incubated at 4 °C with gentle agitation for 48 h to allow phage flocculation. Following incubation, the solution was aliquoted into 50 mL Falcon tubes and centrifuged at 8,000 rpm for 40 min. The resulting precipitate was re-suspended in 2 mL microcentrifuge tubes and further centrifuged at 13,000 rpm for 40 min to concentrate the viral particles. The final phage pellet was re-suspended in 400 µL of SM buffer (HiMedia).

### Library Preparation for Bacteriophage for Next Generation DNA Sequencing

The concentrated bacteriophage was subjected to bacterial DNA extraction using AIT viral Nucleic acid extraction kit (AITbiotech, Singapore). The extracted Phage genomic DNA was first purified using 0.8X AMPure beads. The purified DNA was then tagmented using 0.5µl Amplicon Tagment Mix (ATM) and 2.5µl Tagment DNA Buffer (TD). The tagmentation was performed at 55°C for 10 minutes followed by neutralization adding 1.25µl of Neutralization Tagment Buffer (NT) at room temperature for 5 minutes. The entire tagmented DNA was subjected to indexing using 1.25µl each of i5 and i7 indexes and 3.75µl of Nextera PCR Master Mix (NPM). The thermocycling condition for indexing was as: 72°C for 3 minutes, 95°C for 30s followed by 18 cycles of 95°C for 10s, 55°C for 30s and 72°C for 30s. The final extension was performed at 72°C for 5 minutes. The entire indexed PCR products were purified using 0.7X AMPure beads and quantified using Qubit HS DNA kit. The products’ fragment size was determined using Bioanalyzer (Agilent Technologies, USA) and all the products were normalized to 4nM and pooled together to obtain final library pool of 4nM. The library was spiked with 5% PhiX and denatured using equal volume of 0.2N NaOH. The denatured library was neutralized using Hyb buffer and 10pM of the final library was taken for Sequencing in Illumina MiSeq Reagent V2 kit.

### Bioinformatics analysis for Phage Characterization

The Fastq data were taken, and quality control were carried out with FastP and the filtered reads were assembled with SPAdes genome assembler v4.2 using default parameters, without enabling the “careful” option. The resulting contigs were analyzed with CheckV v1.0.3 [31] for viral contig identification and quality assessment. Contigs classified as complete genomes by CheckV were subsequently annotated and taxonomically classified using PhageScope (https://PhageScope.deepomics.org/). Furthermore, the sequenced whole genome was subjected to antimicrobial and virulence gene analysis with Abricate using CARD and virulence factor database (VFDB) [32–34] (Figure 3).

**Figure 3:**
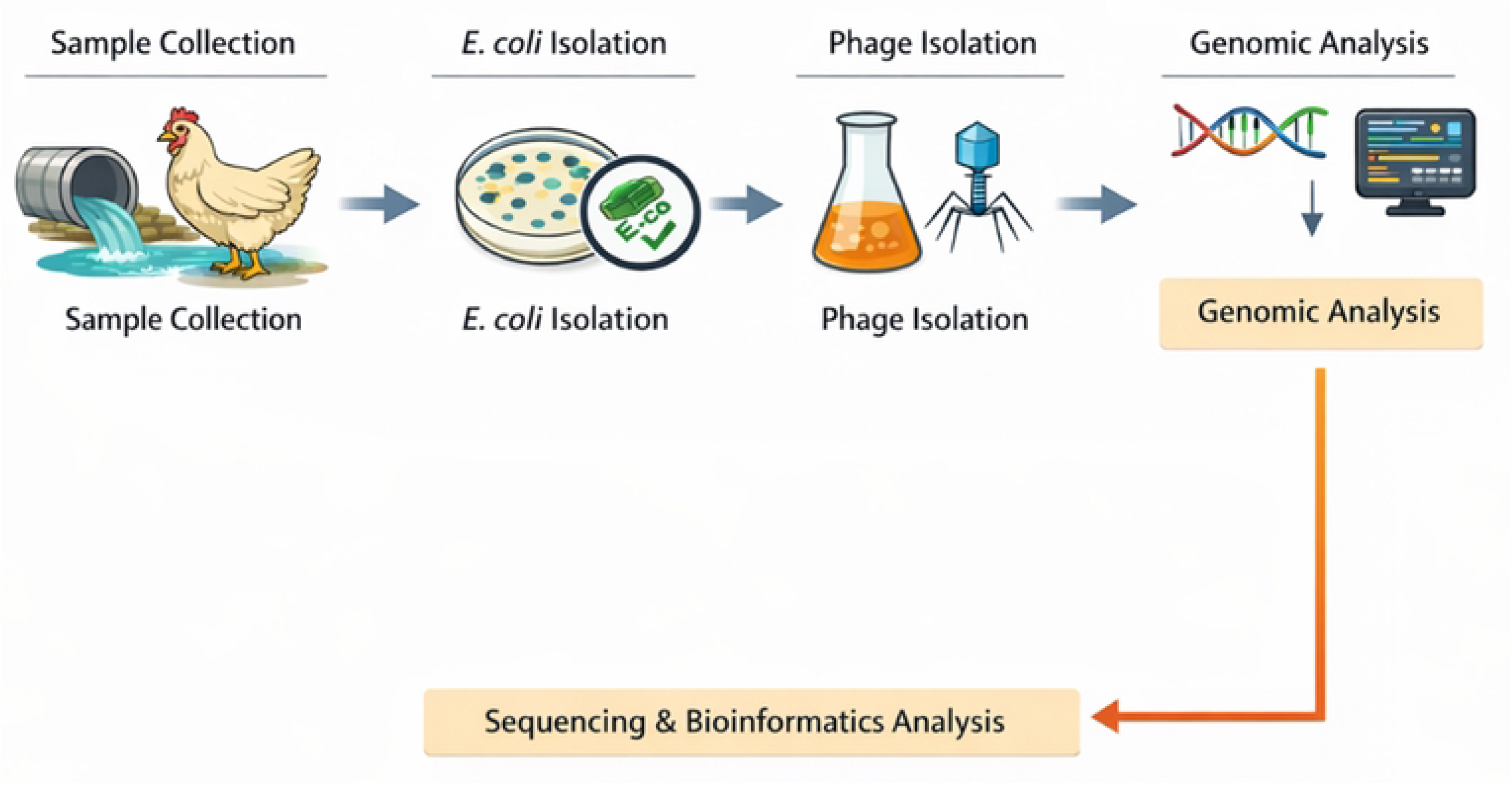
Schematic representation of the study workflow. Biological and environmental samples collected from poultry farms across Nepal were processed for Escherichia coli isolation and confirmation. Confirmed E. coli isolates were used as host strains for bacteriophage enrichment and isolation from sewage samples. Purified phages were concentrated, sequenced, and analyzed using genome assembly, taxonomic classification, and antimicrobial resistance and virulence gene screening pipelines.

## Results

### Prevalence of *E. coli*

*E. coli* was initially identified by traditional culture and biochemical methods and was later confirmed by 16S rRNA sequencing. 19 samples were initially identified as *E. coli* based on phenotypical characteristics i.e., green metallic sheen colonies on EMB agar and yellow/yellow on TSIA, indole positive, MR positive, urease negative and citrate negative. Confirmed 16S sequencing data (PRJNA1393614) verified only 17 isolates as *E. coli*. Two samples were sequenced confirmed to be *Citrobacter* spp, hence were not processed further (Figure 4).

**Figure 4:**
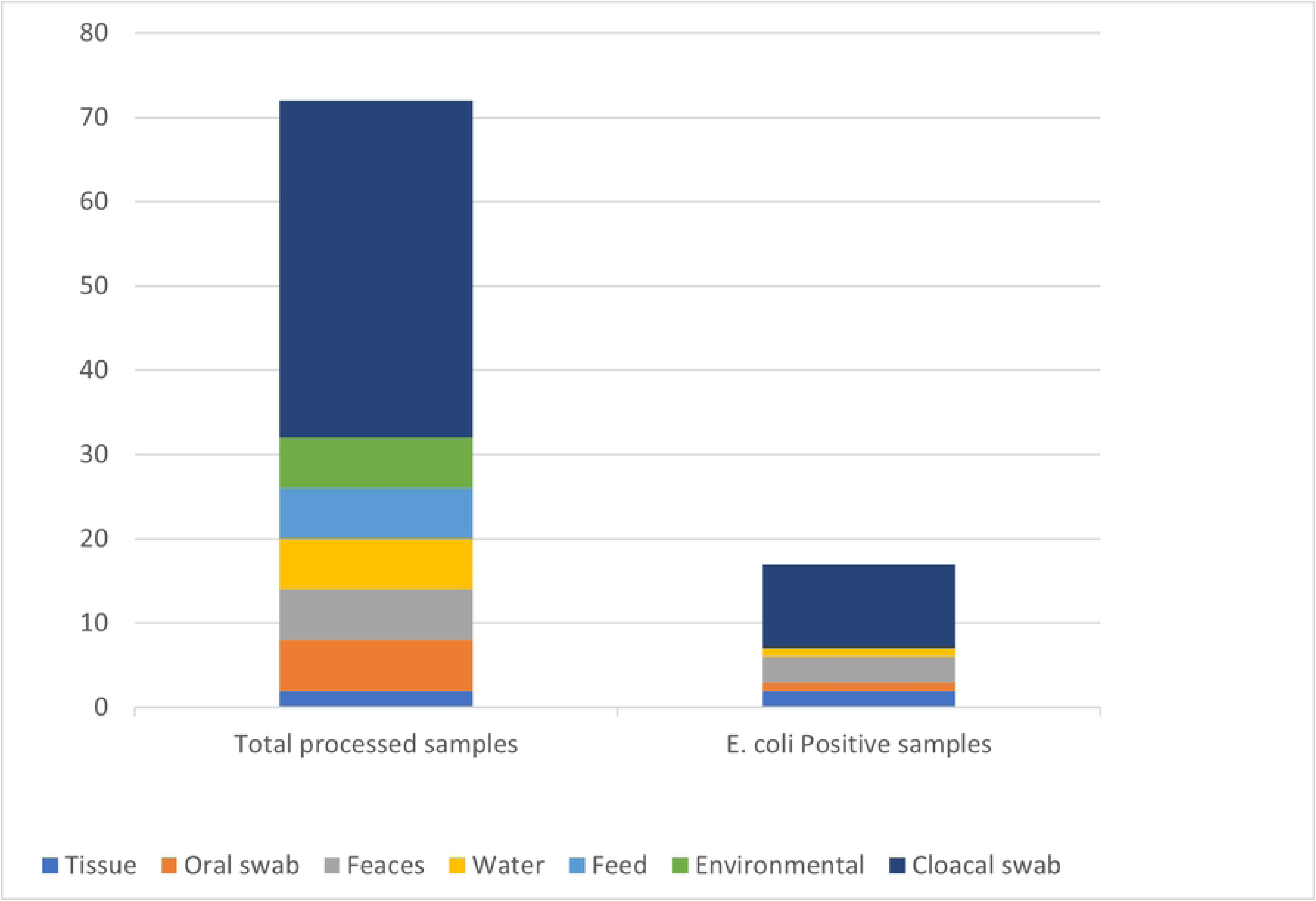
Distribution of poultry and environmental samples and *Escherichia coli* positivity. Stacked bar charts show the total number of samples processed (left) and the number of *E. coli*–positive samples (right) by sample type. Sample categories include tissue, oral swab, feces, water, feed, environmental samples, and cloacal swabs. Cloacal swabs contributed the largest proportion of both total processed samples and *E. coli*–positive detections, highlighting their higher yield for *E. coli* isolation compared with other sample matrices.

Seventeen *E. coli* isolates were obtained from samples collected at nine sites. The majority were from cloacal swabs (10/17, 58.8%), with additional isolates from faeces (3/17, 17.6%), tissue (2/17, 11.8%), farm water (1/17, 5.9%), and an oral swab (1/17, 5.9%). Most isolates were derived from broiler chickens (7/17), followed by mixed local breeds (4/17), layers (3/17), local breeds (2/17), and one Lohmann Brown chicken. While the majority of isolates were collected from healthy birds, five *E. coli* isolates came from sick chickens at the Goldhunga and Chitwan farms. Among these, two tissue samples were derived from necropsy of chickens that had died due to illness.

### Phage Isolation

A total of 18 bacteriophage were isolated. Each isolated phage was initially screened against 17 isolated *E. coli* through double layer agar method. A clear zone over a bacterial lawn was observed due to lytic activity and different phages were concluded based on zone of lysis (Figure 5) and *E. coli* isolates. 18 isolated phages were able to infect 8 *E. coli* isolates. An equal number of phage isolates, specifically six from Balaju sewage, six from Thapathali sewage, and six from Dallu sewage, were generated. Additionally, Phage PG7, PG11, and PG12 were able to infect *E. coli* isolated from sick chicken of Chitwan farm (Table 1). However, none of the phage were able to infect *E. coli* isolated from dead chicken of Goldhunga.

**Figure 5:**
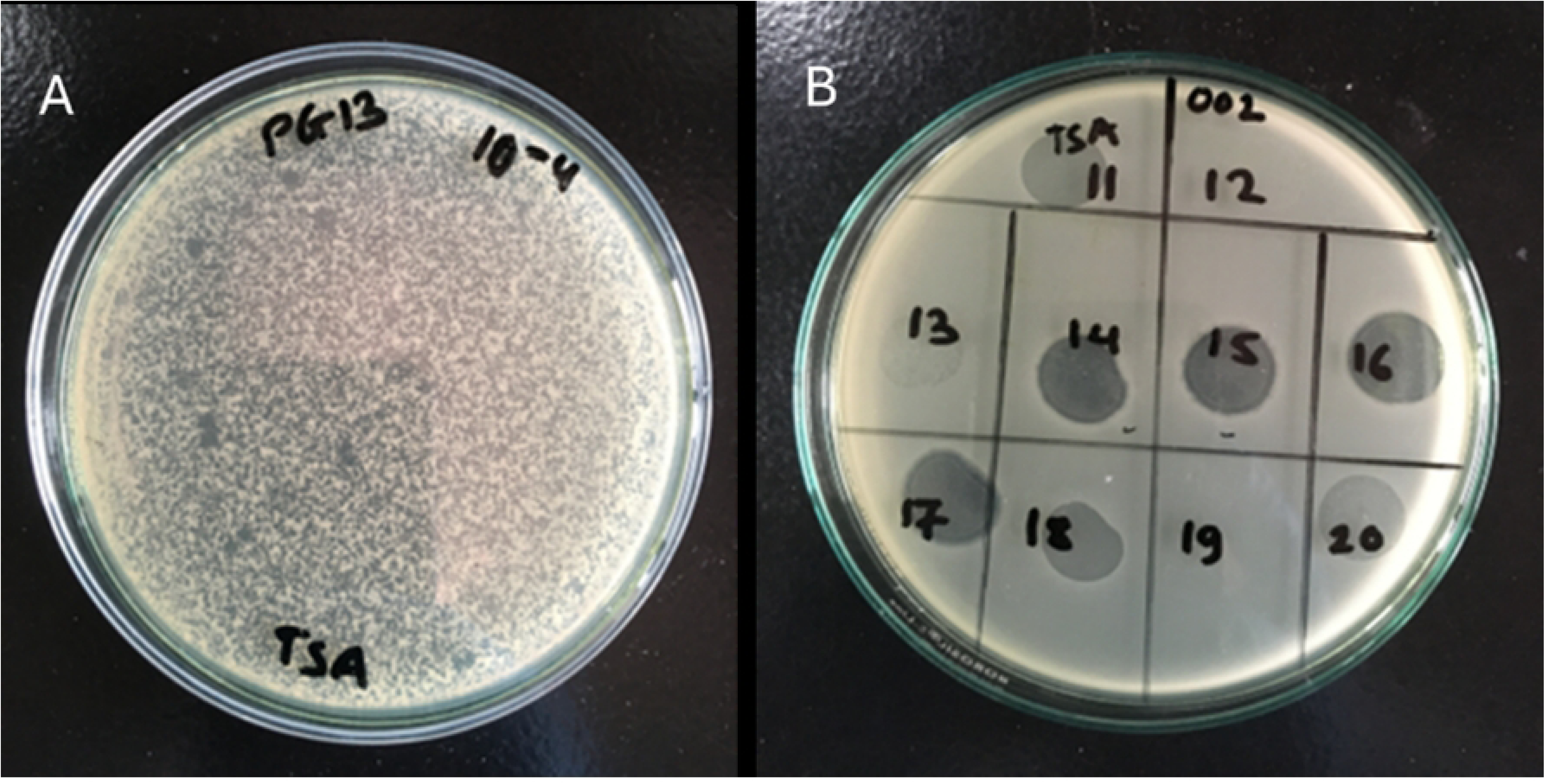
A) Phage plaques (small and big) isolated at 10^-4^ dilution by double layer agar method in TSA medium. B) Cross-infectivity testing of different phages

**Table 1.** List of Phage isolates isolated from different sewage sites against the E. coli hosts used.

| Phage identifier | Lab ID | Life cycle | Source of isolation | Host's (*E. coli*) code |
| --- | --- | --- | --- | --- |
| *E. coli* phage _PG1 | PG1 | Lytic cycle | Sewage from Thapathali | EcI6 |
| *E. coli* phage _PG2 | PG2 | Lytic cycle | Sewage from Balaju | EcI6 |
| *E. coli* phage _PG3 | PG3 | Lytic cycle | Sewage from Thapathali | EcI4 |
| *E. coli* phage _PG4 | PG4 | Lytic cycle | Sewage from Dallu | EcI4 |
| *E. coli* phage _PG5 | PG5 | Lytic cycle | Sewage from Balaju | EcI4 |
| *E. coli* phage _PG6 | PG6 | Lytic cycle | Sewage from Dallu | EcI5 |
| *E. coli* phage _PG7 | PG7 | Lytic cycle | Sewage from Balaju | EcI8 |
| *E. coli* phage _PG9 | PG9 | Lytic cycle | Sewage from Balaju | EcI5 |
| *E. coli* phage _PG10 | PG10 | Lytic cycle | Sewage from Thapathali | EcI5 |
| *E. coli* phage _PG11 | PG11 | Lytic cycle | Sewage from Thapathali | EcI6 |
| *E. coli* phage _PG12 | PG12 | Lytic cycle | Sewage from Thapathali | EcI8 |
| *E. coli* phage _PG13 | PG13 | Lytic cycle | Sewage from Dallu | EcI8 |
| *E. coli* phage _PG14 | PG14 | Lytic cycle | Sewage from Balaju | EcI1 |
| *E. coli* phage _PG15 | PG15 | Lytic cycle | Sewage from Dallu | EcI1 |
| *E. coli* phage _PG16 | PG16 | Lytic cycle | Sewage from Dallu | EcI2 |
| *E. coli* phage _PG17 | PG17 | Lytic cycle | Sewage from Balaju | EcI2 |
| *E. coli* phage _PG18 | PG18 | Lytic cycle | Sewage from Thapathali | EcI9 |
| *E. coli* phage _PG20 | PG20 | Lytic cycle | Sewage from Dallu | EcI17 |

### AMR detection in *E. coli*

Among the eight *E. coli* hosts susceptible to the phages, qPCR screening (Resistomap, Finland) revealed that six carried AMR genes, with a total of 41 different genes detected (Table 2). These genes conferred resistance to seven antibiotic classes: aminoglycosides, rifampin, colistin, MLSB, quinolones, β-lactams, and vancomycin. Notably, the class 1 integron integrase gene intI1_1 was present in all six isolates, intI1_2 in three, and the class 3 integrase intI3 in one.

**Table 2:**
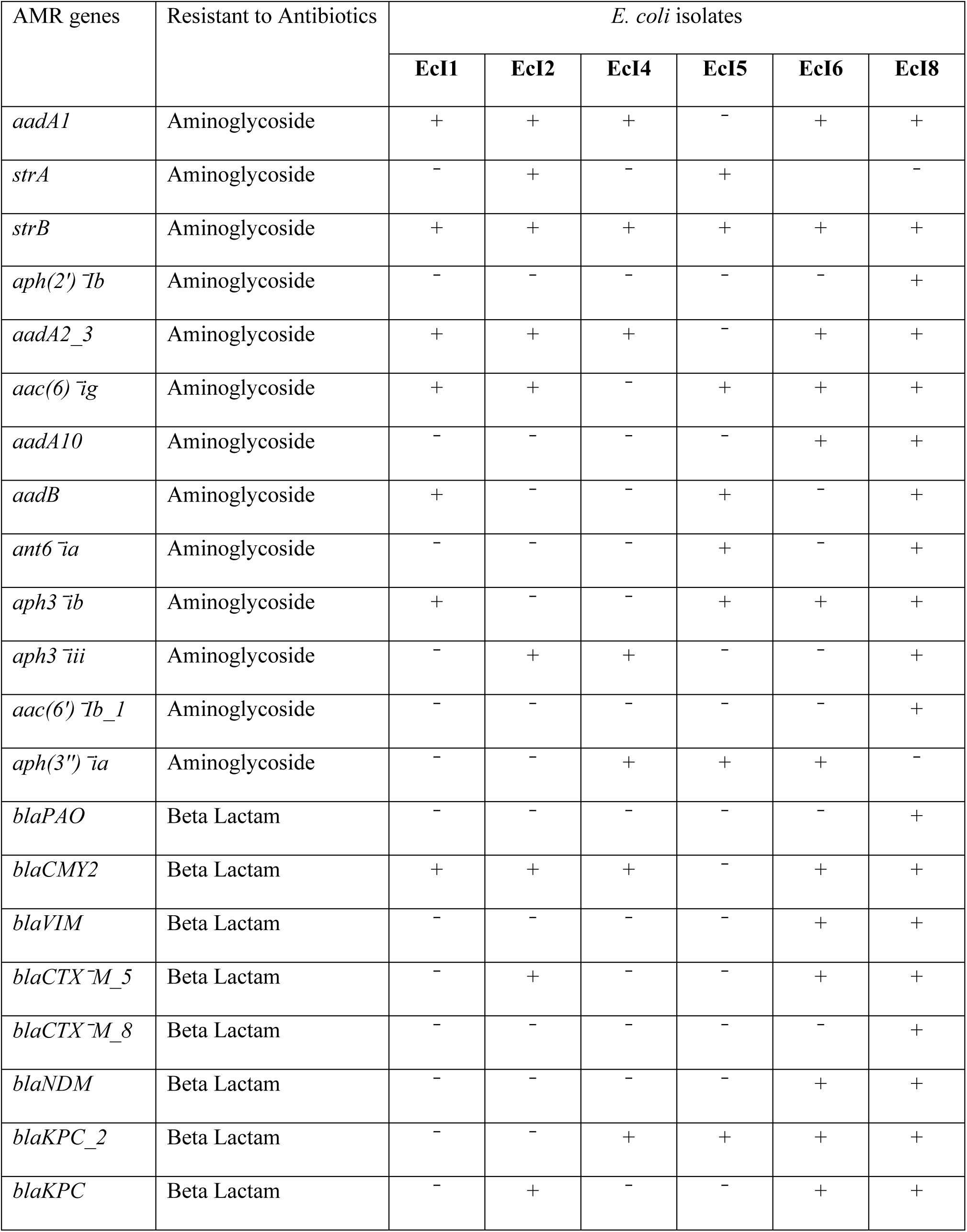
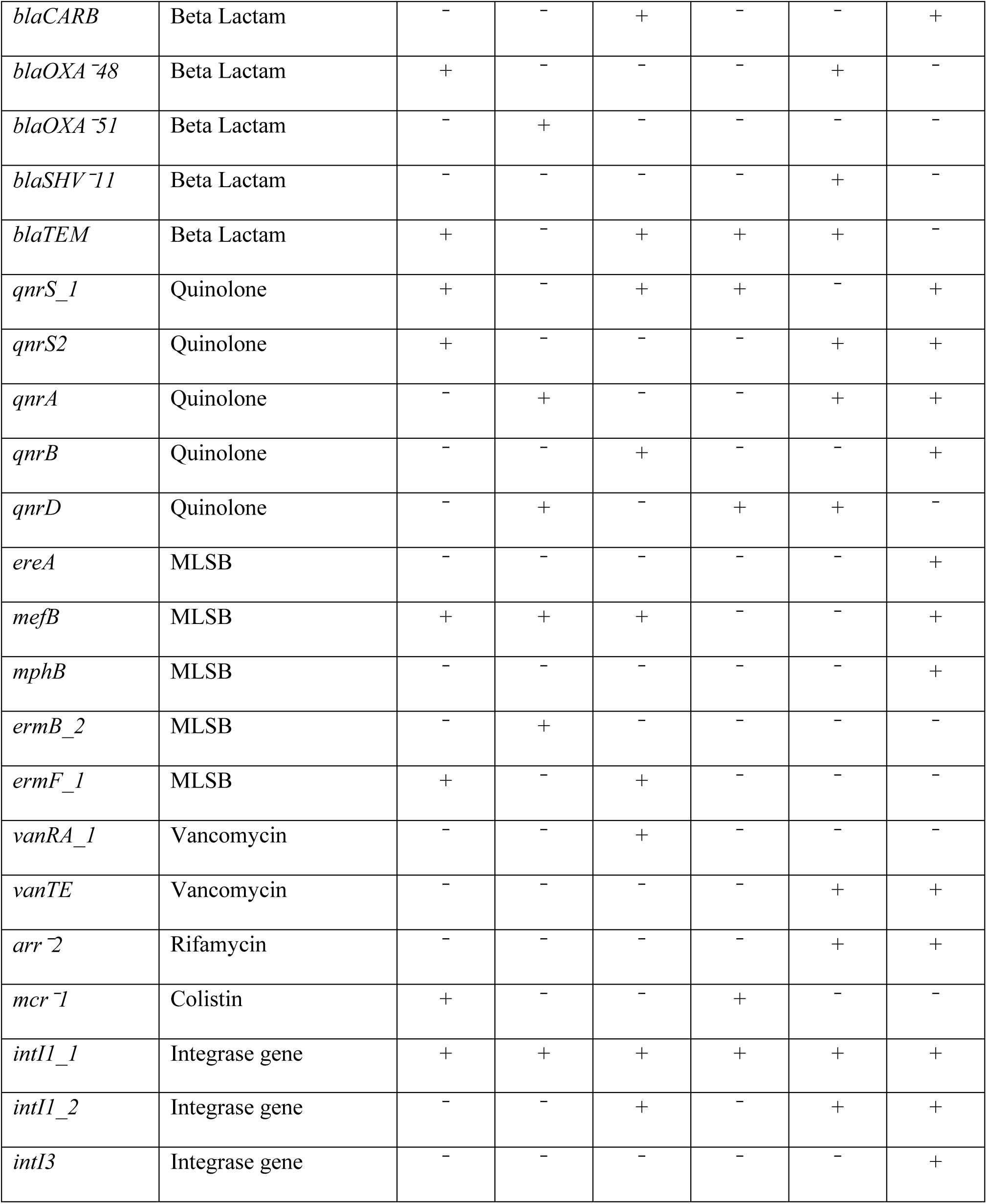
AMR genes detected in six E. coli isolates that are host to different phages using multiplex qPCR assay.

EcI8—the host for phages PG7, PG12, and PG13—showed the highest AMR load, with CT-positive results for 30 genes, spanning 11 aminoglycoside, 10 β-lactam, 4 quinolone, 3 MLSB, and one each of rifamycin and vancomycin resistance, plus two class 1 and one class 3 integron integrase genes.

EcI2, host to PG16 and PG17, carried 14 AMR genes and one class 1 integron gene, including six aminoglycoside, four β-lactam, two MLSB, and two quinolone resistance genes. EcI1, which hosts PG14 and PG15, showed a similar profile with 14 AMR genes (six aminoglycoside, three β-lactam, two MLSB, two quinolone, and one colistin (mcr1)) plus one class 1 integron gene.

EcI6 harbored 21 AMR genes and two class 1 integron genes. In contrast to EcI1, EcI2, and EcI8, it had more β-lactam resistance genes (9) than aminoglycoside (7), lacked MLSB genes, and included three quinolone, one vancomycin, and one rifamycin resistance gene. EcI4 and EcI5 contained 14 and 12 AMR genes respectively, and both possessed class 1 integron integrase genes.

All six isolates also carried carbapenemase genes, conferring resistance to carbapenem antibiotics. In total, six carbapenemase types were detected: blaKPC, blaKPC2, blaOXA-48, blaOXA-51, blaNDM, and blaVIM, with EcI6 containing the highest number (five).

Additionally, EcI1 and EcI5 harbored the mcr-1 gene, which provides resistance to the last-resort antibiotic colistin. Vancomycin resistance genes were found in three isolates: vanRA in EcI4 and vanTE in EcI6 and EcI8. ESBL genes (blaSHV11, blaTEM, blaCTX-M-5, blaCTX-M-8) were also present, with EcI6 carrying three of the four detected ESBL variants.

### Host range testing

Cross infectivity was performed between 18 phages and 8 *E. coli* isolates. Most phages could infect other *E. coli* isolates besides their host *E. coli* except PG5 and PG6. The number of *E. coli* isolates that a particular phage can infect is shown in Figure 6. The most extensive infectivity was observed in PG2, PG4, PG11, PG13, PG16 and PG18 as they were capable of forming plaques in 6 out of 8 *E. coli* isolates, in addition to their respective *E. coli* host. These versatile phages successfully infected *E. coli* that tested positive for antimicrobial resistance (AMR) genes. Specifically, PG2 and PG13 demonstrated infectivity against all 6 AMR-positive *E. coli* isolates, while PG4, PG11, and PG16 were able to infect 5 out of the 6 AMR-positive *E. coli*. Furthermore, almost all other phages, apart from the ones mentioned above, exhibited infectivity towards 5 different *E. coli* isolates (Figure 6).

**Figure 6:**
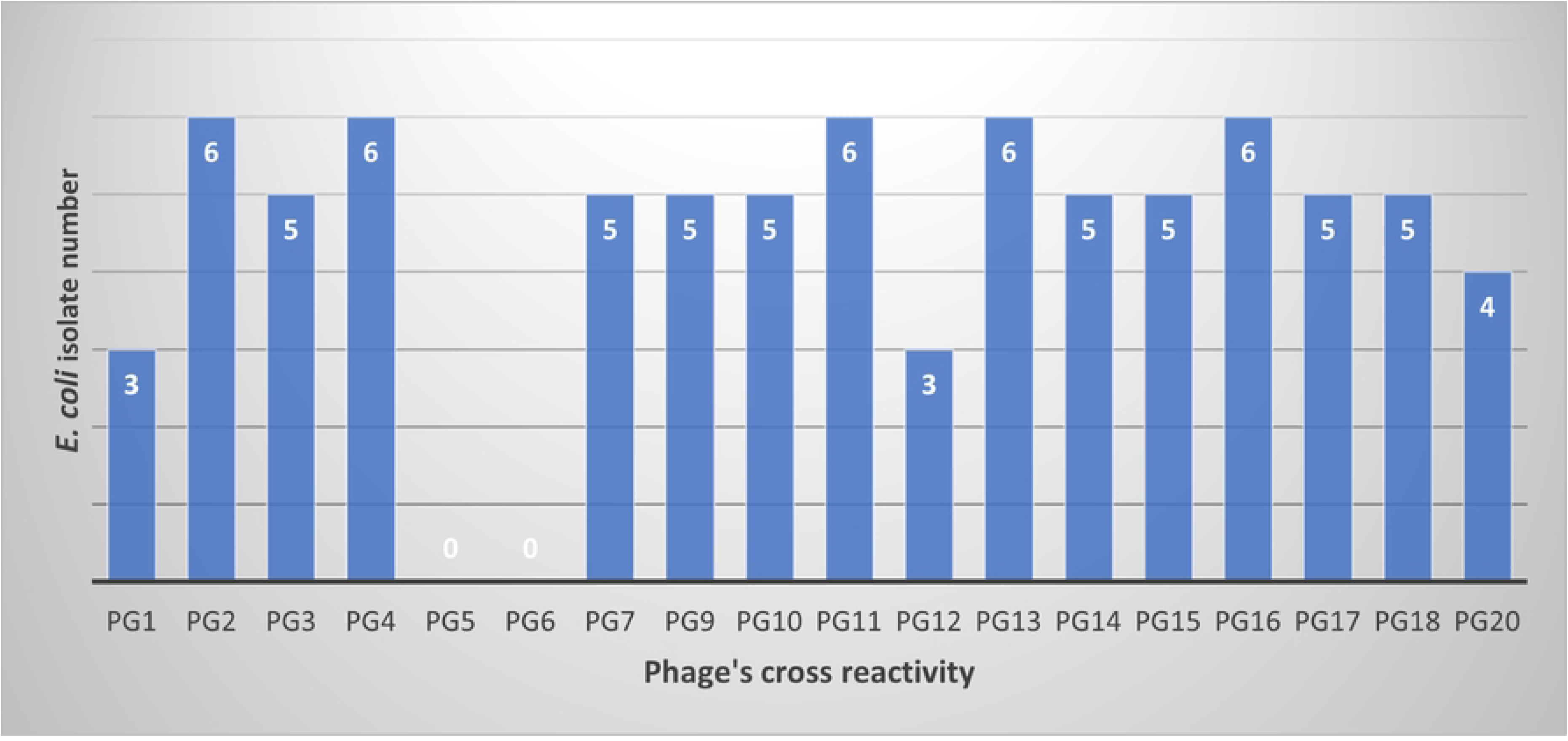
Cross-reactivity of isolated bacteriophages against *Escherichia coli* isolates. The bar chart shows the number of *E. coli* isolates lysed by each bacteriophage (PG1–PG20) during cross-infectivity assays. Phages exhibited variable host ranges, with several phages (e.g., PG2, PG4, PG11, PG13, and PG16) demonstrating broader lytic activity, while others showed limited or no activity (e.g., PG5 and PG6), indicating heterogeneity in phage–host interactions among the tested *E. coli* isolates.

### Phage genomic characterization

Phages were initially isolated from 10⁻⁴ and 10⁻⁶ dilutions, whereas propagated phages were obtained from 10⁻¹⁰ and 10⁻¹² dilutions. Although 18 phages were initially detected, purification through successive passage to obtain uniform plaques resulted in 15 single, pure phages, which were then sequenced on the Illumina MiSeq platform. Passage revealed that some initial isolates contained mixed phages, which were separated into individual phages through repeated passages. For example, PG2 yielded two distinct phages: PG2B5 (large lysis zone) and PG2S3 (small lysis zone), while PG5 as separated into PG5_1 and PG5. Similarly, PG6 and PG8 were split into PG6_1, PG6, PG8, and PG8_1.

### Genomic and functional characterization of phages

Sequencing results indicated that seven phages had insufficient or low coverage: PG4 produced only a partial genome (<10% coverage), PG2_S3 had 3.58× coverage, and PG5_1, PG6, PG6_1, PG8, and PG8_1 were also excluded due to inadequate coverage. Consequently, only six phages had sufficient coverage for genomic characterization (Figure 7). All sequencing data have been submitted to the NCBI SRA database under accession number PRJNA1393614.

**Figure 7:**
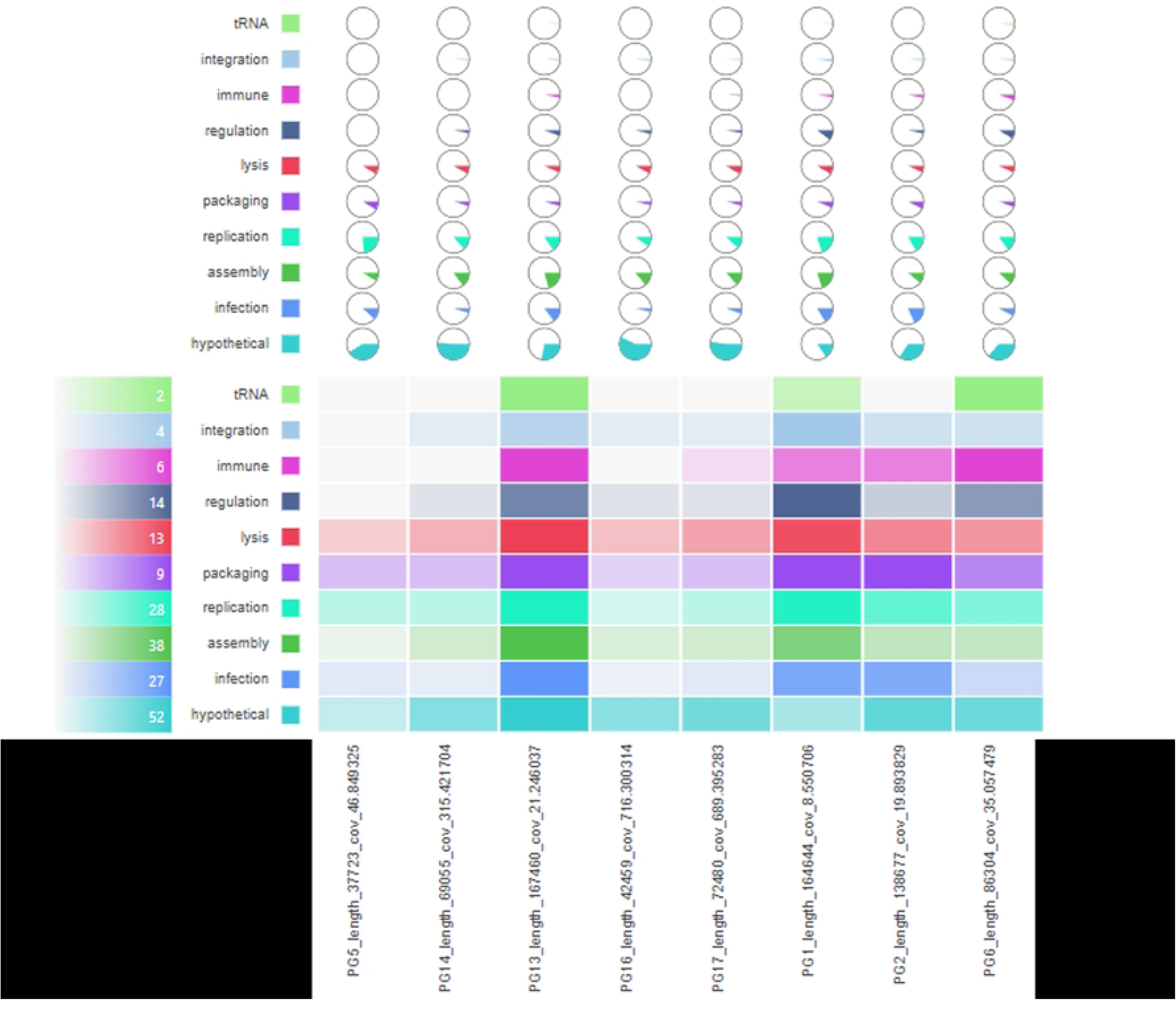
Functional gene composition of representative *Escherichia coli* bacteriophage genomes. The figure summarizes the distribution of predicted open reading frames (ORFs) across major functional categories for selected bacteriophages. Circular glyphs illustrate the proportional representation of functional modules, while the heatmap below shows the absolute number of genes assigned to each category, including tRNA, integration, immune evasion, regulation, lysis, packaging, replication, assembly, infection, and hypothetical proteins. Phage genomes are ordered by size, highlighting conserved core functions involved in replication, assembly, and lysis, alongside substantial proportions of hypothetical proteins, reflecting genetic diversity and incomplete functional annotation among the isolated phages.

Initial sequencing revealed that all eight purified phages belonged to four families within the Caudovirales order (**Table 3**). Three phages (PG14, PG16, PG17) were assigned to the Podoviridae family, Gamaleyavirus genus. Within Myoviridae, PG2B5 was classified as a Vequintavirus (138.58 Kbp) and showed similarity to Escherichia phage vB_EcoM-ECP26, while PG13B6 and PG6 were Tequatroviruses (167.361 Kbp and 86.304 Kbp), similar to Escherichia phage UGKSEcP2. PG5 was identified as a member of the Chaseviridae family. Genome analysis confirmed that none of the sequenced phages carried antimicrobial resistance or virulence genes capable of transduction [35] or conferring pathogenicity to *E. coli*.

**Table 3:** Classification of purified phages based on sequencing results and similarity to known phage. N/D= not determined.

| **Phage** | **Genus** | **Subfamily** | **Family** | **Order** | **Genome size**<br>**(Kbp)** | **Comparable to this**<br>**phage** | **Accession no.** | **Similarity**<br>**%** |
| --- | --- | --- | --- | --- | --- | --- | --- | --- |
| **PG1** | Krischvirus | — | Straboviridae | Caudovirales | N/D | Escherichia phage<br>FL15 | PP400778.1 | 96.64 |
| **PG2B5** | Vequentavirus | Vequentavirinae | Myoviridae | Caudovirales | 138.58 | Escherichia phage<br>vB_EcoM-ECP26 | MK883717.1 | 99.22 |
| **PG5** | — | Cleopatravirinae | Chaseviridae | Caudovirales | 37.723 | Escherichia phage<br>Ecp_YSF | OR327751.1 | 96.83 |
| **PG6** | Tequatrovirus | Tevenvirinae | Myoviridae | Caudovirales | 86.304 | — | — | — |
| **PG13B6** | Tequatrovirus | Tevenvirinae | Myoviridae | Caudovirales | N/D | Escherichia phage<br>UGKSEcP2 | OV876900.1 | 99.05 |
| **PG14** | Gamaleyavirus | Enquatrovirinae | Podoviridae | Caudovirales | 69.055 | Escherichia phage<br>vB_EcoP-ZQ2 | NC_070874 | 93.07 |
| **PG16** | Gamaleyavirus | Enquatrovirinae | Podoviridae | Caudovirales | 42.459 | — | — | — |
| **PG17** | Gamaleyavirus | Enquatrovirinae | Podoviridae | Caudovirales | 72.381 | Escherichia phage<br>vB_EcoP-ZQ2 | NC_070874 | 93.71 |

Genome analysis revealed that all eight sequenced phages (Figure 7, Supplementary Figures 1-8) carried genes encoding RZ-like spanins. PG1 and PG5 also contained multiple lysis-related genes, such as T4 holin [36], Gene 25-like lysozyme. PG6 encoded enzymes with lytic transglycosylase [37] and cysteine-type peptidase functions. PG14 and PG17 included genes associated with metallopeptidase-mediated lysis. PG13 and PG17 harbored Gene 25-like lysozyme, lytic transglycosylase [37], and bacteriophage T-type holin genes, all contributing to bacterial cell wall breakdown. Finally, PG16 encoded holin involved in cell lysis [38].

## Discussion

*E. coli* causes colibacillosis, a common poultry disease causing significant mortality and economic losses, but antibiotic treatments such as neomycin are increasingly ineffective due to antimicrobial resistance and are no longer recommended [39][40][41][42]. Currently, prevention relies mainly on hygiene and sanitation, yet effective treatment and control strategies remain inadequate [43][44][45]. In this study, we developed an alternative biological approach by rapidly isolating and genomically characterizing naturally occurring bacteriophages and evaluating their efficacy against *E. coli* in poultry. This strategy has the potential to address both colibacillosis and antimicrobial resistance in the field.

Most *E. coli* isolates were obtained from cloacal swabs (58.82%, 10/17), followed by fecal samples (17.65%, 3/17). As a common member of the chicken gut microbiome, its presence in these samples was expected [46][47]. However, isolates from tissues such as the gizzard and proventriculus (EcI10 and EcI11) were recovered from sick birds, likely associated with colibacillosis [48]. Additionally, *E. coli* detected in farm water suggested fecal contamination and biosafety lapses, while its presence in an oral swab indicated possible colonization.

E. coli isolates (EcI8 and EcI9) from Chitwan farms were recovered from chickens infected with Influenza A virus, suggesting secondary colibacillosis following viral infection [49]. Given the higher susceptibility of broiler chickens to diseases [50], most of the *E. coli* isolates were isolated from broilers, followed by mixed local varieties, which are more vulnerable due to lower antibiotic supplementation in their feed [51]. Suspected colibacillosis cases were observed exclusively in broilers.

Extensive antibiotic use in poultry, particularly in broilers, has promoted multidrug-resistant (MDR) E. coli with high antimicrobial resistance (AMR) gene burdens [52][44]. In this study, isolate EcI8 from broiler feces in Chitwan carried the highest number of resistance genes across multiple antibiotic classes including 11 aminoglycoside, 4 quinolone, 3 MLSB, 1 rifamycin, 1 vancomycin, and 9 β-lactam resistance genes (Table 2), likely facilitated by the presence of class 1 and 3 integrons [53][54]. Similar MDR profiles were observed in isolates from other farms (EcI6 EcI1, EcI2, and EcI4) (Table 1), reflecting differences in management practices, breed, and antibiotic exposure [55][56][57]. Given the rapid turnover of broilers and their role in the human food chain, these findings raise serious concerns about zoonotic transmission of livestock-associated AMR to humans [58][59][60][61]. Strong associations have already been documented between antimicrobial use in food animals and the prevalence of resistant bacteria in both animals and humans [62][63] [64].

From three sewage samples, we isolated 18 distinct bacteriophages that lysed eight highly resistant poultry-derived E. coli isolates, including the most resistant strain, EcI8 (Table 1 and Table 2). Cross-infectivity assays confirmed that the phages were unique and non-redundant (Figure 2; Table 2 and **Table 3**). Several phages showed broad host ranges: PG13 (Tequatrovirus), PG2 (Vequintavirus), and PG16 (Gamaleyavirus) each infected seven *E. coli* isolates, consistent with the association between larger genome size and wider host range [65][66]. Importantly, phages from genera previously used in animal and human phage therapy trials were recovered, i.e., Tequatrovirus (PG13) [67], Phapecoctavirus (PG7) [68][69], Vequintavirus (PG2, PG2B5) [70], and Gamaleyavirus (PG14, PG16, PG17) (46)(47). Notably, PG7—closely related to the APEC-targeting phage ESCO5—lysed MDR strains and lacked detectable toxin, virulence, or resistance genes, supporting its therapeutic potential [68][69].

Phage susceptibility varied markedly even among closely related isolates from the same farm (e.g., EcI5 vs. EcI6; EcI1 vs. EcI2), supporting phage host-range profiling as an additional strain-typing tool [71][72][73]. Phages active against *E. coli* from clinically affected chickens likely suffering from secondary bacterial infection following primary viral disease [74]) in Chitwan-belonged to genera previously used in veterinary phage [68] [75][76]. In contrast, no lytic phages were recovered for EcI11 and EcI12 from deceased birds, highlighting variability in phage susceptibility among poultry *E. coli* strains. Overall, sewage-derived bacteriophages—particularly Tequatrovirus, Phapecoctavirus, Vequintavirus, and Gamaleyavirus—show promise as antibiotic-free interventions against MDR *E. coli* in poultry. These phages could be formulated into farm-specific cocktails and administered via feed or drinking water, enabling scalable integration into existing poultry production systems [77].

Colibacillosis, a common and severe poultry disease caused by E. coli, has become an increasing concern due to rising antibiotic resistance [78][68]. The identification of both broad- and narrow-host-range phages capable of effectively lysing *E. coli* is therefore highly encouraging. Sewage samples from Balaju, Dallu, and Thapathali contained a diverse range of *E. coli*-infecting phages, highlighting sewage as a rich reservoir of therapeutic candidates [79]. Rapid isolation, characterization, and use of phages against new and re-emerging bacterial pathogens like *E. coli* is critical for making phage treatment viable and as effective as antibiotics. Although phages have considerable promise, their use has been limited, which can be addressed in part by quick isolation and characterization. Introducing phage isolation and characterization dynamics might improve Colibacillosis treatment and prevention strategies, helping the poultry business and lessening the impact of antibiotic resistance [80][81][82].

In addition to their use as whole-phage biocontrol agents against MDR *E. coli*, the eight fully sequenced phages encoded diverse cell-wall-degrading enzymes with potential as standalone antimicrobials. All phages carried RZ-like spanin genes, and several contained additional lysis proteins, including T4 holin [36], Gene 25-like lysozyme, lytic transglycosylases [35], cysteine-type peptidases, metallopeptidases, and bacteriophage T-type holins—enzymes collectively responsible for efficient disruption of the peptidoglycan layer and outer membrane during the phage lytic cycle. Importantly, none of these phages carried antimicrobial resistance or virulence genes capable of transduction [35], ensuring that the lysis genes are not linked to undesirable genetic elements. These phage-derived lytic enzymes—particularly endolysins, spanins, and hydrolases from PG13, PG17, PG6, PG14, and PG16—could be developed as recombinant enzybiotics, offering advantages such as non-replicative action, broader activity in combination, and simpler regulatory pathways [83–85]. Delivered via feed or water, alone or alongside phage cocktails, they represent a complementary strategy for controlling antibiotic-resistant *E. coli* in poultry.

While this study demonstrates the feasibility of rapid phage isolation and genomic characterization as an alternative to antibiotics for controlling MDR *E. coli* in poultry, several limitations should be noted. The small sample size—17 poultry isolates from a limited number of farms in three Nepalese districts and phages from only three sewage sites—restricts generalizability. The rapid pipeline emphasized speed over extensive characterization, leaving key parameters such as phage stability, growth kinetics, morphology, and delivery optimization unassessed. All analyses were conducted in vitro, and phage efficacy, dosing, microbiome interactions, and host immune responses remain to be validated in animal models. In addition, the absence of comparisons with existing phage therapies or other antibiotic alternatives limits translational interpretation. Nevertheless, the results highlight clear directions for future work, including broader sampling, in vivo trials, phage cocktail optimization, and field validation to advance phage-based control of avian colibacillosis and antimicrobial resistance.

## Conclusion

In this study, we demonstrated that multidrug-resistant *E. coli* is prevalent in poultry, particularly in commercial broiler chickens, with isolates carrying high burdens of antimicrobial resistance genes and posing potential zoonotic risks. These findings underscore the urgent need for effective antibiotic-free control strategies.

Using a rapid isolation and genomic characterization pipeline, we recovered a diverse collection of naturally occurring bacteriophages from sewage that were active against highly resistant poultry-derived *E. coli* strains. Several phages exhibited broad host ranges and belonged to genera previously validated in veterinary and human phage therapy, while lacking detectable virulence or resistance genes, supporting their therapeutic safety and relevance. Variability in phage susceptibility among closely related *E. coli* isolates further highlighted the value of phage host-range profiling for strain-level targeting and cocktail design.

Beyond whole-phage applications, the identified phages encoded multiple lytic enzymes with strong potential as recombinant enzybiotics, offering complementary or alternative antimicrobial approaches with simpler regulatory pathways. Although limited by sample size and in vitro validation, this study establishes proof-of-concept that rapid phage discovery and genomic screening can generate safe, effective candidates for controlling multidrug-resistant *E. coli* in poultry. Collectively, these findings support sewage-derived bacteriophages and phage-derived enzymes as promising, scalable tools to mitigate colibacillosis and reduce reliance on antibiotics, while providing a clear framework for future in vivo testing and field-level implementation.

## Acknowledgement

We sincerely thank all field and laboratory team members at BIOVAC Nepal and the Center for Molecular Dynamics Nepal for their dedicated contributions to this study. We are also grateful to Resistomap, Finland, for supporting the qPCR-based screening of antimicrobial resistance (AMR). Finally, we thank our collaborative team at Phulping Farm Pvt. Ltd. for their invaluable assistance with field logistics.

**Supplementary Figures (1-8): Whole genome sequence annotations of phage isolates**

**Supplementary Figure 1**: Whole genome sequence annotation of phage isolate **PG1** with gene name on the label and differently coloured gene as per the function.

**Supplementary Figure 2**: Whole genome sequence annotation of phage isolate **PG2** with gene name on the label and differently coloured gene as per the function.

**Supplementary Figure 3**: Whole genome sequence annotation of phage isolate **PG5** with gene name on the label and differently coloured gene as per the function.

**Supplementary Figure 4**: Whole genome sequence annotation of phage isolate **PG6** with gene name on the label and differently coloured gene as per the function.

**Supplementary Figure 5**: Whole genome sequence annotation of phage isolate **PG13** with gene name on the label and differently coloured gene as per the function.

**Supplementary Figure 6**: Whole genome sequence annotation of phage isolate **PG14** with gene name on the label and differently coloured gene as per the function.

**Supplementary Figure 7**: Whole genome sequence annotation of phage isolate **PG16** with gene name on the label and differently coloured gene as per the function.

**Supplementary Figure 8**: Whole genome sequence annotation of phage isolate **PG17** with gene name on the label and differently coloured gene as per the function.

